# Reconstructing heterogeneous pathogen interactions from co-occurrence data via statistical network inference

**DOI:** 10.1101/2021.11.15.468692

**Authors:** Irene Man, Elisa Benincà, Mirjam E. Kretzschmar, Johannes A. Bogaards

## Abstract

Infectious diseases often involve multiple pathogen species or multiple strains of the same pathogen. As such, knowledge of how different pathogen species or pathogen strains interact is key to understand and predict the outcome of interventions that target only a single pathogen or subset of strains involved in disease. While population-level data have been used to infer pathogen strain interactions, most previously used inference methods only consider uniform interactions between all strains, or focus on marginal interactions between pairs of strains (without correction for indirect interactions through other strains). Here, we evaluate whether statistical network inference could be useful for reconstructing heterogeneous interaction networks from cross-sectional surveys tracking co-occurrence of multi-strain pathogens. To this end, we applied a suite of network models to data simulating endemic infection states of pathogen strains. Satisfactory performance was demonstrated by unbiased estimation of interaction parameters for large sample size. Accurate reconstruction of networks may require regularization or penalizing for sample size. Of note, performance deteriorated in the presence of host heterogeneity, but this could be overcome by correcting for individual-level risk factors. Our work demonstrates how statistical network inference could prove useful for detecting pathogen interactions and may have implications beyond epidemiology.

## Introduction

The rapid expansion of sequencing technologies over the last decades has drastically increased our ability to detect within-host pathogen diversity. As a result, it is recognised that infectious diseases often involve multiple pathogen species. For example, diseases may arise from bacterial and viral co-infections [1, 2], or co-infections of multiple strains of the same pathogen species, e.g. serotypes of *Streptococcus pneumoniae* [3], genotypes of the human papillomavirus [4], or clones of *Plasmodium falciparum* [5]. It is also increasingly understood that networks of microbial interactions may determine the outcome of preventive or therapeutic interventions (e.g. vaccination, antibiotics, probiotics) in unexpected ways [6–8]. For infectious agents, these interactions between pathogens or pathogen strains can take various forms: infection by one strain can modify susceptibility to subsequent infection by others, and the simultaneous presence of multiple strains can affect the duration, infectiousness, as well as severity of infection [9, 10]. Consequently, pathogen interactions may have far-reaching clinical, epidemiological, and eco-evolutionary implications [11–13].

The acknowledgement that pathogen strain interaction is a crucial factor shaping the ecology for a wide number of infectious agents has spurred the development of epidemiological models where hosts can be infected by different pathogens or different strains of the same pathogen [14]. However, inference of interaction parameters in these models is notoriously challenging, especially when pathogen strain interaction is to be extracted from population-level data. The availability of large-scale cross-sectional surveys, detecting the presence of multiple infectious strains simultaneously, has given rise to a wide range of methods for detecting interactions from co-occurrence data [15–20]. These methods rely on identifying deviation from statistical independence, either in the number of pathogen strains occurring in individual hosts, pairwise co-occurrence between any two strains, or the exact combination of co-occurrence.

Clearly, the validity of such statistical methods for detecting pathogen interactions hinges on the assumption that non-interaction implies statistical independence, but statistical associations can be driven by factors other than biological interactions [21]. This point has been made repeatedly by community ecologists, concerned with the possibility to detect interdependence among species from co-occurrence patterns across habitats [22], and has been reiterated by epidemiologists [23]. A common criticism of using statistical associations is that hosts or habitats are not identical, and this may favour the presence of one or another (or both) species irrespective of biological interactions. Opportunely, epidemiological studies on human pathogens typically offer rich meta-data, making it possible (in principle) to correct for host heterogeneity. Moreover, when studying different strains of the same pathogen, one might assume that the same hosts are more likely than others to become infected with all strains, justifying correction for host heterogeneity by host-specific covariates in statistical models [23].

So far, many existing methods for detecting multi-strain pathogen interactions are restricted to uniform (or homogeneous) interactions across strains (i.e. all strains under consideration interact with one another in the same manner), or focused on marginal interactions between pairs of pathogen strains. Marginal estimation of pairwise interactions does not necessarily control for indirect interactions through other strains and may therefore be prone to bias, especially when strains interact in a heterogeneous manner. Network inference methods relying on conditional dependency between the co-occurrence of multiple strains might be more suited for detecting heterogeneous pathogen strain interactions (e.g. interactions that are different between each pair of strains), but are not widely considered. Moreover, it is not yet well studied how the performance of these methods might be affected by the presence of host heterogeneity.

The purpose of this paper is two-fold. First, we explore the possibility of detecting heterogeneous pathogen interactions from cross-sectional survey data using statistical network inference methods. For this, we evaluate the performance of a suite of network models in reconstructing simulated networks of pathogen strain interactions for a range of epidemiological and network settings. Second, we investigate the robustness of these methods to settings with host heterogeneity that is relevant for pathogen spread. In this work, we focus on reconstruction of interaction network of pathogen strains, but the methodology is applicable to cross-sectional surveys tracking co-occurrence of potentially interacting pathogen species and may have implications beyond epidemiology.

## Results

In order to evaluate the performance of various network inference methods, we simulated random interaction networks with up to 10 pathogen strains, which describe whether and to what extent the strains interact in an epidemiological model. Within each network, the presence of interaction between each pair of strains *i* and *j* was established with connection probability *σ*. Given that interaction between a pair was established, the strength of interaction, indicated by the value of *x*_*ij*_, was drawn uniformly from the range [−*θ, θ*], where *θ* is a positive number. Values of *x*_*ij*_ in the range [−*θ*, 0) and (0, *θ*] correspond to competitive and mutualistic interactions, respectively, and the greater the value diverges from zero, the stronger the interaction becomes.

Each network generated as such was then used to parametrize a Susceptible-Infected-Susceptible (SIS) epidemiological model [8]. In the model, exp(*x*_*ij*_) was used to as the hazard ratio that determines how the presence of strain *i* in a host alters the host’s hazards to acquire or clear strain *j*, and vice versa. See Material and methods for the exact description of how parameter *x*_*ij*_ is utilized in the parametrization of the epidemiological model. After parametrization, the steady state prevalence of the epidemiological model was obtained and used to sample random cross-sectional datasets with the number of observations being 100, 1.000, 10.000, or 100.000. These datasets were then used to reconstruct the interaction networks. See S1 Fig for a schematic representation of the entire data generation process.

The set of network inference methods that we considered were the Ising model, graphical modelling approaches, and generalized estimating equations (GEE) [24–26]. For the Ising model and GEE, model selection was done based lasso (least absolute shrinkage and selection operator) and Wald tests with false discovery rate control, respectively. Graphical model selection was performed either in a forward or a backward fashion (i.e. by iteratively growing or shrinking networks, respectively), using either AIC or BIC as selection criterion.

In the base-case setting, we considered 500 random networks with 5 strains, each generated with connection probability *σ* = 0.25 and a range of interaction parameters of [−*θ, θ*] = [− log 3, log 3]. In the corresponding epidemiological model, we considered interaction in acquisition only (i.e. no interaction in clearance), a population without host heterogeneity, and type-specific basic reproduction numbers *R*_0,*i*_ in the range [1.5, 2].

In the base-case setting, visual inspection showed satisfactory concordance between the estimated networks and true networks. This is illustrated in Fig 1, which also shows some differences between network inference methods. In order to compare the different methods systematically, we assessed various performance measures, including sensitivity and specificity for indicating the presence (or absence) of pairwise interactions, i.e. the proportions of truly present (or truly absent) interactions that are correctly identified in the reconstructed networks. In addition, we assessed the F1-score, which summarizes sensitivity and positive predictive value (PPV, i.e. the proportion of identified interactions that are truly present) in the form of their harmonic mean.

**Fig 1.**
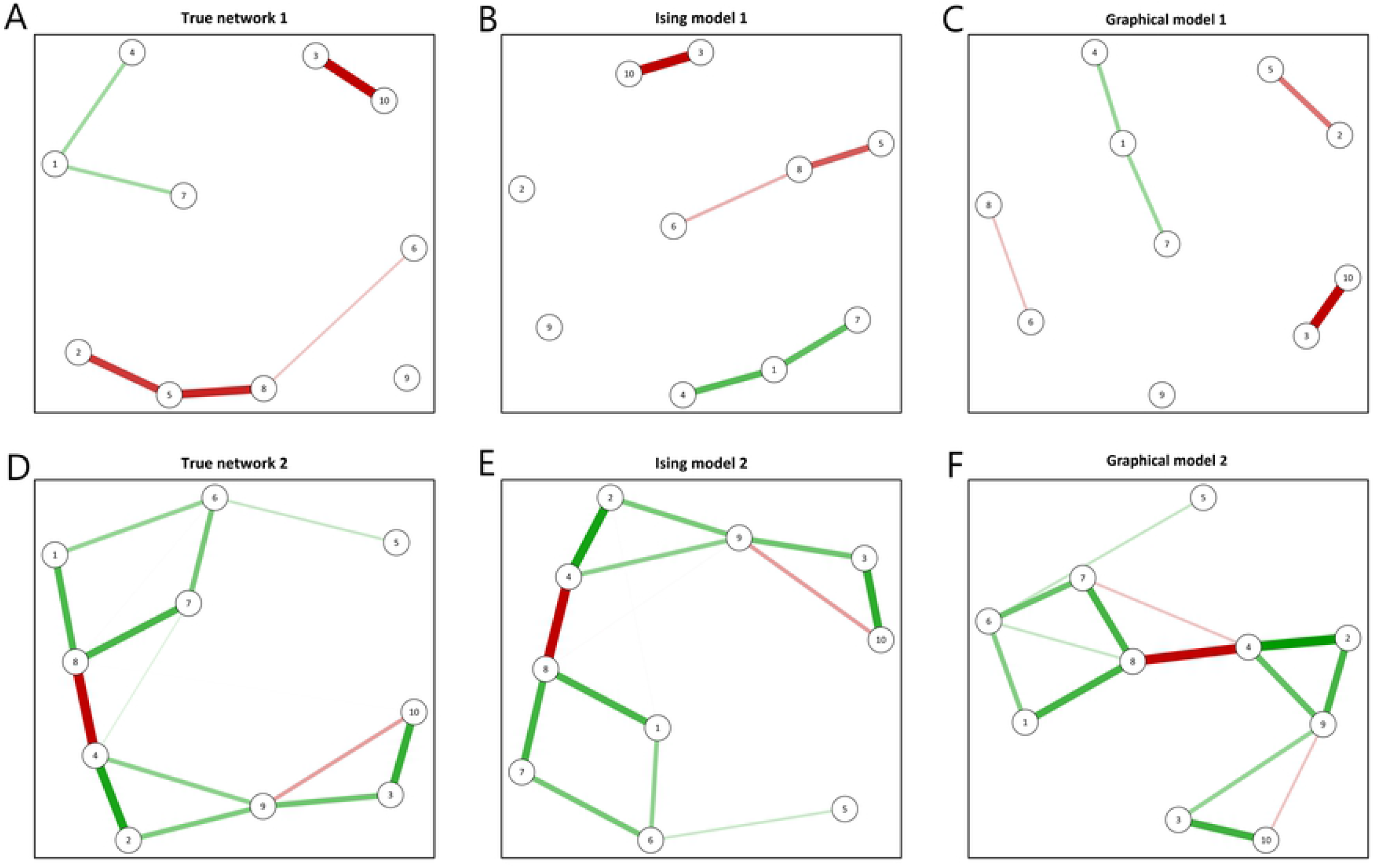
Examples of networks with *n* = 10 strains in the simulation study. Two random networks generated with connection probability *σ* = 0.25 and interaction strengths *x*_*ij*_ drawn from [−*θ, θ*] = [− log 3, log 3] (A and D); Estimated networks from co-occurrence in 100,000 observations according to the regularized Ising model (B and E); and according to a graphical model using backward model selection by BIC (C and F). Strength and type of interactions (mutualistic versus competitive) are indicated by the thickness and colour (green versus red) of edges, respectively.

### Trade-off between sensitivity and specificity

Network reconstruction improved with the number of observations according to almost all assessed performance measures for all considered methods (Fig 2A-D, Table 1). Interactions were recovered with almost 100% sensitivity at 100,000 observations (Fig 2A). Optimal specificity was attained already at small sample size in graphical modelling approaches with BIC-based, whereas specificity remained at a more or less constant suboptimal level in graphical modelling approaches with AIC-based selection (Fig 2D). GEE was good at small sample size, but was the only method considered with slightly deteriorating performance with increasing sample size notwithstanding correction for multiple testing (Fig 2B-D). Overall, methods with less optimal specificity showed better sensitivity, showcasing trade-off between sensitivity and specificity. Finally, the Ising model and the graphical modelling approaches with BIC-based selection were able to achieve relatively high values of sensitivity and specificity at high sample size, which resulted in near-optimal F1-scores. (Fig 2B,C).

**Fig 2.**
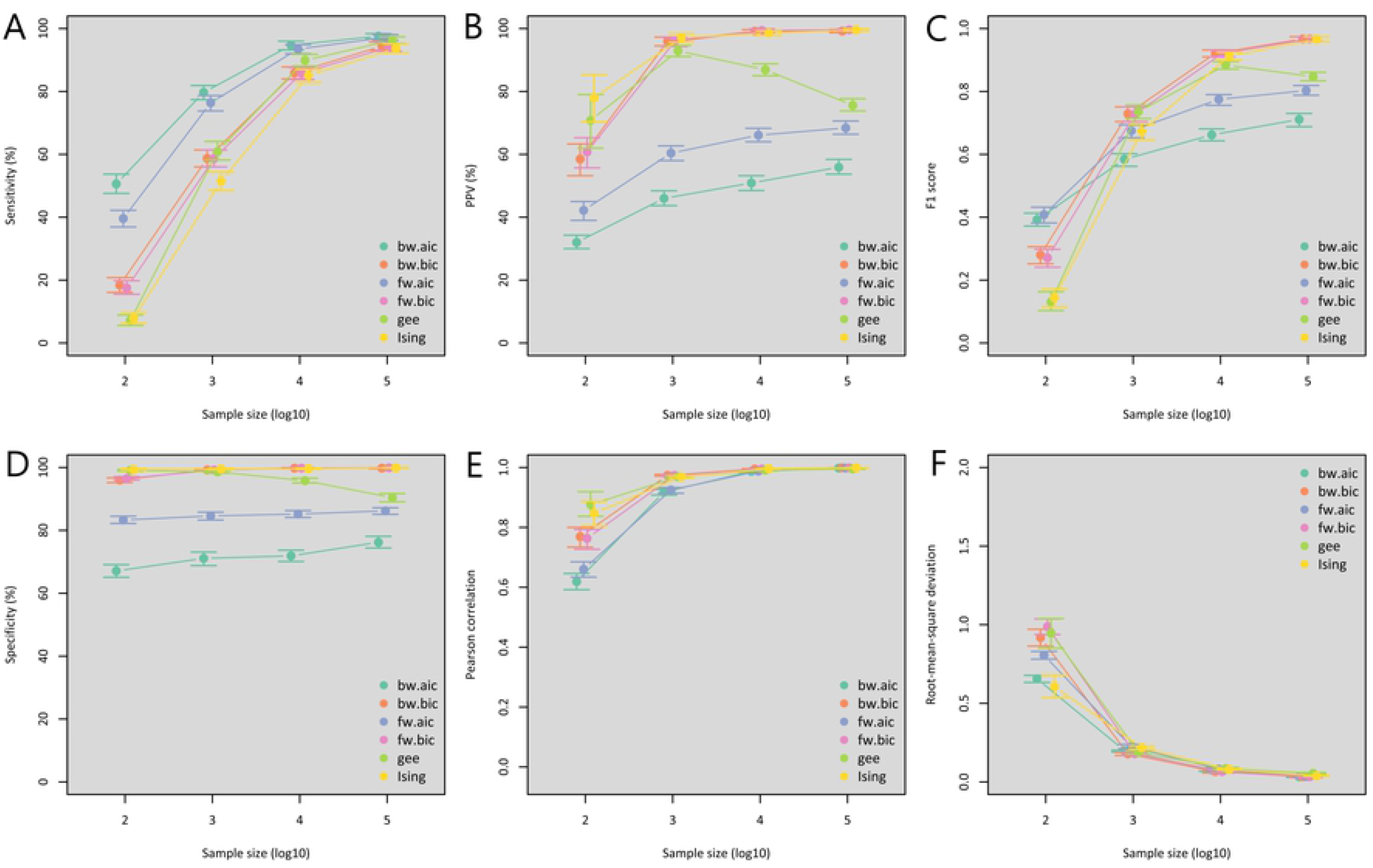
Performance measures of statistical network inference in the base-case analysis. Several measures have been calculated to assess the performance of the statistical network inference as a function of the sample size for the base-case analysis. A) Sensitivity; B) Positive predictive value (PPV); C) F1 score; D) Specificity; E) Pearson correlation; F) Root-mean-square deviation. In the base-case analysis, interactions between strains were generated with connection probability *σ* = 0.25, interaction strengths *x*_*ij*_ indicating interaction in acquisition drawn from [−log 3, log 3], and basic reproduction numbers randomly drawn from [1.5, 2]. All methods were evaluated at sample sizes of 100, 1,000, 10,000 and 100,000 observations, but x-axis coordinates are slightly jittered to improve visualization. Abbreviations: bw.aic (turquoise): graphical model with backward AIC selection; bw.bic (red): graphical model with backward BIC selection; fw.aic (purple): graphical model with forward AIC selection; fw.bic (pink): graphical model with forward BIC selection; gee (green): generalized estimating equations; Ising (yellow): Ising model.

**Table 1.** Performance measures of statistical network inference in the base-case analysis. In the base-case analysis, interactions between strains were generated with connection probability *σ* = 0.25, interaction strengths *x*_*ij*_ indicating interaction in acquisition drawn from [−log 3, log 3], and basic reproduction numbers randomly drawn from [1.5, 2]. All methods were evaluated at sample sizes of 100, 1,000, 10,000 and 100,000 observations. Abbreviations: GEE, generalized estimating equations; PPV, positive predictive value; RMSE, root-mean-square error.

| Method | Sample size § | Specificity (%) ± | Sensitivity (%) ± | PPV (%) ± | F1-score †± | Pearson's r *± | RMSE ± |
| --- | --- | --- | --- | --- | --- | --- | --- |
| *Ising model* | 100,000 | 99.9 (99.8; 100.) | 93.7 (92.3; 95.1) | 99.6 (99.2; 99.9) | .966 (.958; .974) | .999 (.998; .999) | .038 (.036; .041) |
|  | 10,000 | 99.6 (99.4; 99.8) | 85.1 (82.8; 87.0) | 98.6 (97.8; 99.3) | .913 (.899; .925) | .996 (.995; .996) | .079 (.075; .084) |
|  | 1,000 | 99.5 (99.3; 99.7) | 51.5 (48.5; 54.3) | 97.1 (95.6; 98.4) | .673 (.646; .697) | .969 (.965; .973) | .218 (.208; .229) |
|  | 100 | 99.3 (99.1; 99.6) | 7.9 (6.3; 9.7) | 78.0 (70.3; 85.4) | .144 (.115; .172) | .847 (.805; .887) | .606 (.537; .679) |
| *GEE* | 100,000 | 90.4 (89.1; 91.7) | 96.3 (95.3; 97.4) | 75.6 (73.8; 77.7) | .847 (.834; .861) | .995 (.994; .996) | .055 (.050; .060) |
|  | 10,000 | 95.8 (95.1; 96.6) | 89.9 (88.1; 91.8) | 86.9 (85.0; 88.8) | .884 (.871; .895) | .991 (.989; .992) | .085 (.077; .093) |
|  | 1,000 | 98.6 (98.2; 99.0) | 60.9 (58.2; 64.1) | 92.9 (91.1; 94.6) | .736 (.714; .757) | .969 (.966; .973) | .189 (.177; .199) |
|  | 100 | 99.1 (98.8; 99.4) | 7.2 (5.6; 8.1) | 70.8 (62.0; 79.0) | .131 (.103; .163) | .876 (.837; .919) | .946 (.851; 1.04) |
| *Graphical model*<br>*with backward*<br>*AIC selection* | 100,000 | 76.2 (74.4; 78.1) | 97.6 (96.7; 98.4) | 55.9 (53.7; 58.4) | .711 (.688; .730) | .998 (.997; .998) | .033 (.029; .037) |
|  | 10,000 | 71.9 (70.1; 73.7) | 94.7 (93.2; 96.0) | 50.9 (48.5; 53.2) | .662 (.643; .681) | .987 (.983; .989) | .075 (.067; .085) |
|  | 1,000 | 71.1 (68.8; 73.1) | 79.7 (77.4; 81.9) | 46.0 (43.7; 48.4) | .584 (.562; .602) | .918 (.908; .927) | .200 (.190; .202) |
|  | 100 | 67.1 (65.1; 69.1) | 50.6 (47.6; 53.7) | 32.0 (30.0; 34.3) | .392 (.372; .413) | .619 (.592; .646) | .656 (.633; .678) |
| *Graphical model*<br>*with backward*<br>*BIC selection* | 100,000 | 99.8 (99.6; 99.9) | 94.4 (93.1; 95.7) | 99.2 (98.7; 99.7) | .967 (.960; .974) | .999 (.998; .999) | .033 (.027; .040) |
|  | 10,000 | 99.8 (99.6; 99.9) | 85.9 (84.0; 87.8) | 99.2 (98.5; 99.7) | .921 (.910; .932) | .995 (.994; .996) | .066 (.057; .077) |
|  | 1,000 | 99.2 (99.0; 99.5) | 58.7 (56.0; 61.4) | 95.9 (94.5; 97.3) | .729 (.704; .751) | .974 (.971; .977) | .178 (.168; .187) |
|  | 100 | 96.0 (95.2; 96.7) | 18.4 (16.1; 20.8) | 58.5 (53.2; 63.3) | .280 (.252; .306) | .769 (.734; .800) | .918 (.864; .971) |
| *Graphical model*<br>*with forward*<br>*AIC selection* | 100,000 | 86.2 (85.1; 87.2) | 97.2 (96.3; 98.1) | 68.4 (66.4; 70.6) | .803 (.788; .819) | .998 (.997; .998) | .034 (.030; .040) |
|  | 10,000 | 85.2 (84.1; 86.3) | 93.5 (92.0; 94.9) | 66.1 (64.0; 68.3) | .775 (.756; .790) | .988 (.985; .991) | .081 (.071; .093) |
|  | 1,000 | 84.6 (83.3; 85.8) | 76.4 (73.8; 78.7) | 60.4 (58.0; 62.7) | .675 (.653; .694) | .925 (.914; .933) | .224 (.212; .239) |
|  | 100 | 83.3 (82.2; 84.5) | 39.6 (36.9; 42.2) | 42.2 (39.0; 45.0) | .408 (.382; .432) | .660 (.634; .685) | .806 (.781; .830) |
| *Graphical model*<br>*with forward*<br>*BIC selection* | 100,000 | 99.9 (99.8; 100.) | 94.1 (92.8; 95.3) | 99.6 (99.1; 99.9) | .967 (.960; .974) | .999 (.998; .999) | .033 (.027; .040) |
|  | 10,000 | 99.8 (99.7; 100.) | 85.8 (83.8; 87.6) | 99.4 (98.7; 99.9) | .921 (.910; .932) | .995 (.994; .996) | .066 (.057; .077) |
|  | 1,000 | 99.2 (98.9; 99.5) | 58.7 (56.0; 61.4) | 95.8 (94.3; 97.2) | .728 (.704; .751) | .974 (.970; .977) | .179 (.169; .189) |
|  | 100 | 96.5 (96.0; 97.1) | 17.5 (15.5; 19.8) | 60.8 (55.7; 65.3) | .271 (.241; .298) | .763 (.727; .793) | .987 (.938; 1.04) |

§ Number of sampled individuals
± 95% confidence intervals (bootstrap percentile estimates) in brackets
† F1-score denotes the harmonic mean between sensitivity (also termed recall) and PPV (also termed precision)
\* Linear correlation coefficient between generated and estimated interaction strengths (non-zero estimates only)

### Unbiased estimation of the strength of interaction

In addition, we assessed the quantitative agreement between the estimated and the true interaction parameter *x*_*ij*_, which indicates the strength of interaction, based on Pearson correlation and root-mean-square deviation. Average Pearson correlations between the true and estimated interactions were high for all methods with 1,000 observations or more (Fig 2E). Under the smallest sample size involving only 100 observations, fewer interactions were recovered and the estimated interaction strength deviated somewhat from the true parameters, but even in this case the average correlation between true and estimated interaction strength remained substantial, especially under GEE or regularized Ising estimation (Table 1). Moreover, while correlations between true and estimated interactions reached a plateau above 1,000 observations, accuracy continued to improve with increasing sample size, as verified by steadily decreasing root-mean-square deviations for all methods considered, showcasing asymptotically unbiased estimation (Fig 2F).

### Performance under alternative epidemiological settings and network configurations

In a series of sensitivity analyses, we modified the base-case setting to alternative epidemiological settings or network configurations: inclusion of interaction in clearance, lower basic reproduction numbers *R*_0,*i*_ ∈ [1, 1.5], stronger interactions *x*_*ij*_ ∈ [− log 10, log 10], higher connection probabilities *σ* = 0.5, larger networks by increasing the number of strains to 10, or larger networks by connecting two sub-networks of 5 strains.

Network inference was not affected by including interactions in clearance relative to the base-case setting. In this setting, F1-scores remained the same for all methods considered (compare pink dots to black lines in Fig 3). This implies that the combined effect of interactions in acquisition and clearance was well captured by the estimated interaction parameters. Specifically, the ratio of the interaction parameters in acquisition and clearance was well estimated.

**Fig 3.**
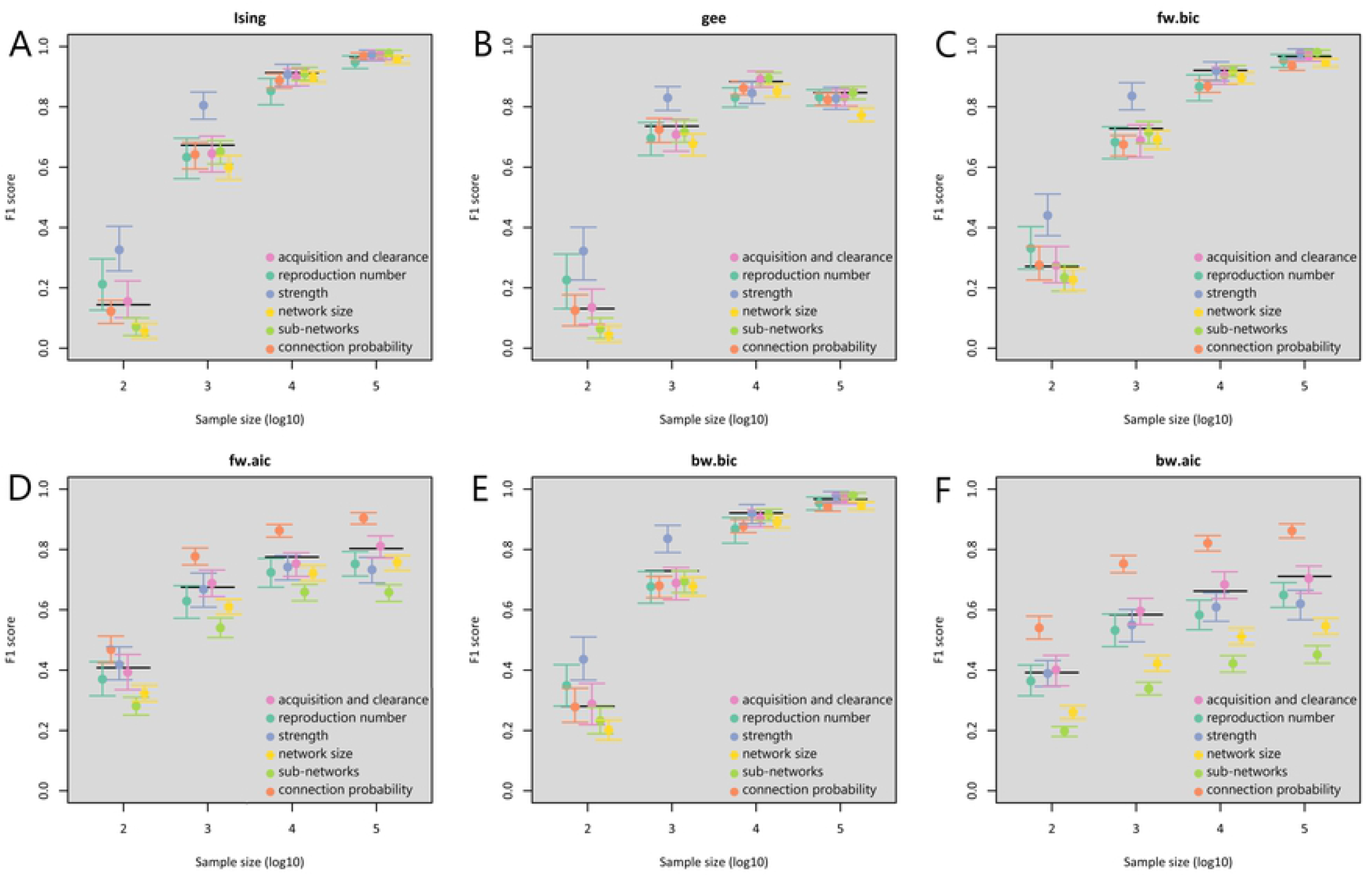
F1-scores of statistical network inference methods in the sensitivity analyses. The performance of different network inference methods expressed by F1-score as a function of the sample size. A) Ising model; B) generalized estimating equations; and graphical models with C) forward BIC selection; D) forward AIC selection; E) backward BIC selection; F) backward AIC selection. The alternative settings considered in the sensitivity analyses were: acquisition and clearance (pink): interaction strengths *x*_*ij*_ including both interactions in acquisition and clearance; reproduction number (turquoise): lower basic reproduction numbers from the range [1, 1.5] instead of [1.5, 2]; strength (purple): strong interaction strengths *x*_*ij*_ drawn from [−log 10, log 10] instead of [−log 3, log 3]; network size (yellow): larger networks with 10 strains; sub-networks (green): larger networks with 10 strains created by collating sub-networks of 5 strains; connection probability (red): higher connection probability *σ* being 0.5 instead of 0.25. Performance of the base-case analysis is given by black horizontal lines.

Looking across all alternative settings, regularized Ising, GEE estimation and BIC-graphical modelling approaches maintained the high performance of the base-case analysis when the sample size was 10,000 or larger (Fig 3A,B,C,E). For these methods, when the sample size was 1,000 or smaller, performance stayed more or less the same with higher connection probability or lower reproduction numbers (compare red and turquoise dots to black lines), improved with stronger interactions (compare purple dots to black lines) but slightly deteriorated when considering larger networks of 10 strains (compare green and yellow dots to black lines).

As for the AIC-based graphical modelling approaches, the performance diverged more from that of the base-case setting (Fig 3D,F). The performance improved substantially under higher connection probabilities (compare red dots to black lines). However, it suffered also more when networks were larger (compare green and yellow dots to black lines). These patterns are linked to the high sensitivity and poor specificity of the AIC-based approaches (as shown in the base-case analysis).

### Robustness to host heterogeneity

Finally, we tested the robustness of the network inference methods to host heterogeneity in contact rate relevant to pathogen spread. The host population was divided into two interconnected sub-populations: one with high and one with low contact rate. Network inference strongly deteriorated in the presence of host heterogeneity as compared to the base-case analysis when heterogeneity was not corrected for (compare turquoise dots to black lines in Fig 4, compare D to A in Fig 5). For all methods considered, F1-scores did not improve with increasing sample size. Similarly, root-mean-square deviation remained high at large sample size (S1 Table). These findings show a consistent bias towards positive associations in uncorrected analyses (S2 Fig).

**Fig 4.**
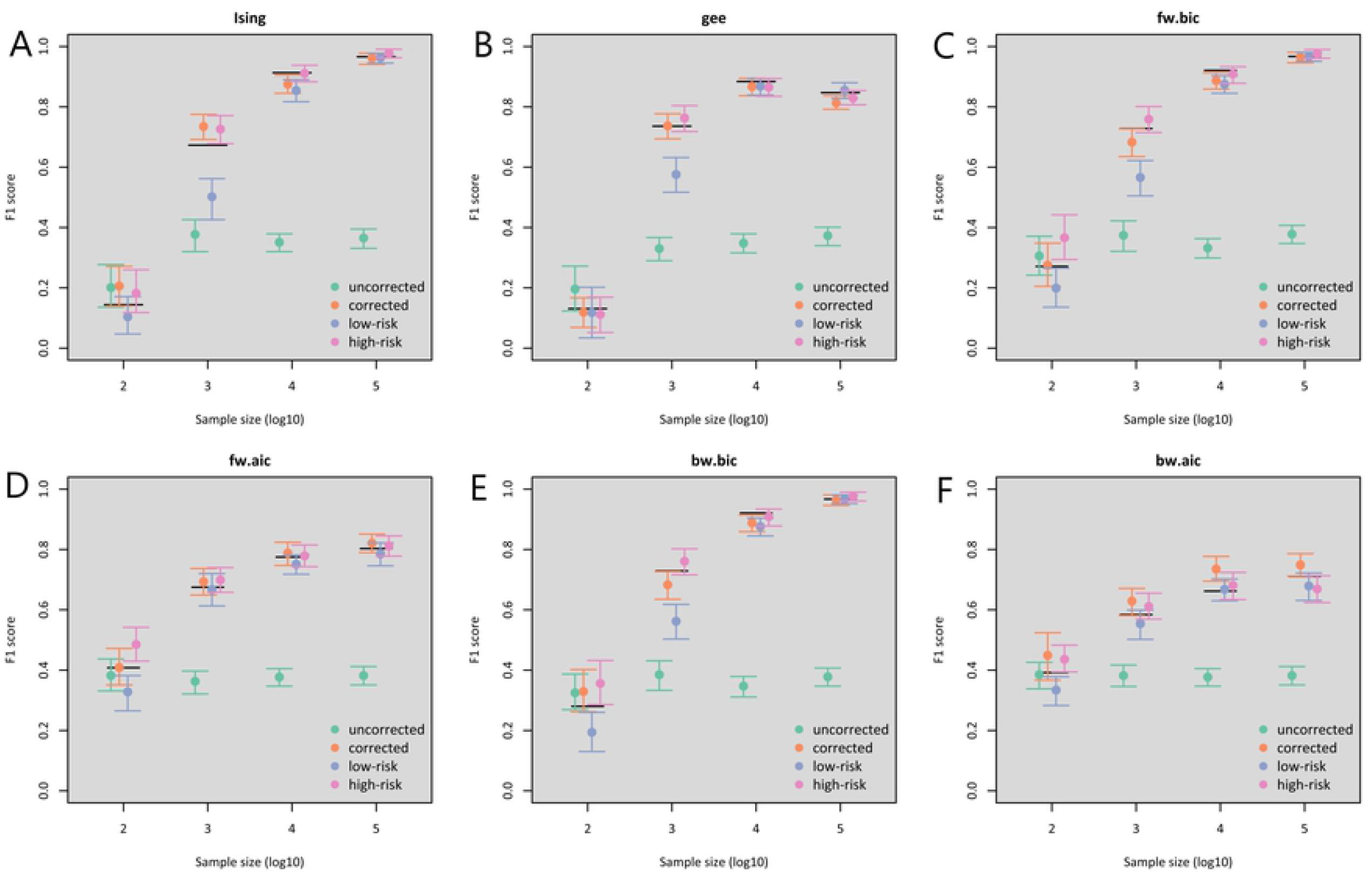
F1-scores of statistical network inference under host heterogeneity. The performance of the different inference methods evaluated under the setting with host heterogeneity as a function of the sample size. A) Ising model; B) generalized estimating equations; and graphical models with C) forward BIC selection; D) forward AIC selection; E) backward BIC selection; F) backward AIC selection. Similar epidemiological models are used as in Fig 2, but with two sub-populations of hosts. Average contact rate is the same as in the base-case analysis, but 80% of hosts is assumed to have below-average contacts and 20% above-average contacts (coefficient of variation: 80%). Mixing between sub-populations occurred pseudo-assortatively (assortivity fraction: 50%). Performance is investigated in following ways: uncorrected: based on representatively sampled individuals from the total population without correction for contact rate; corrected: idem but with correction for contact rate; low-risk: (stratified) analysis on individuals sampled from sub-population with low contact rate only; high-risk: (stratified) analysis on individuals sampled from sub-population with high contact rate only.

**Fig 5.**
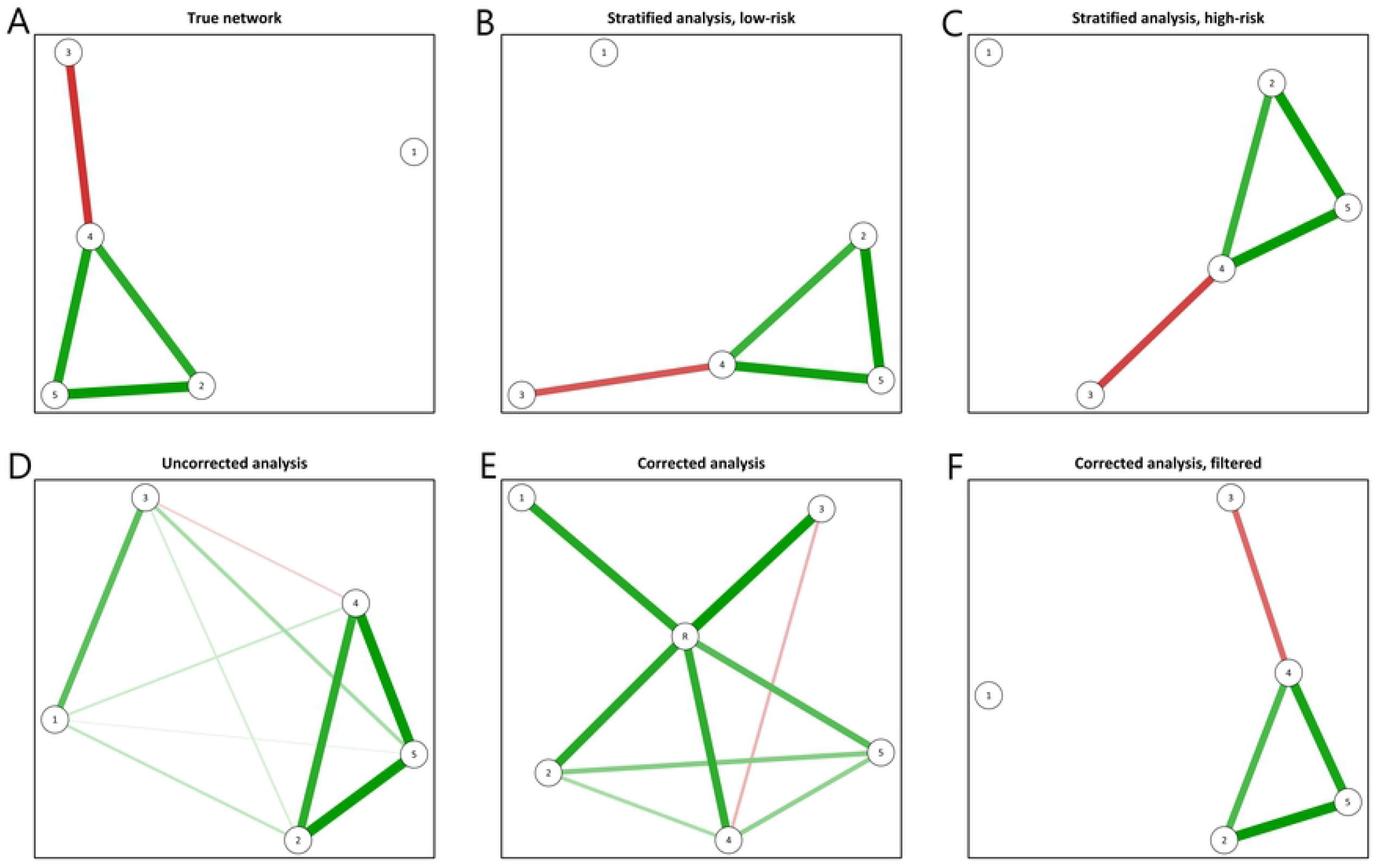
An example of network reconstruction under host heterogeneity. The true random network was generated with connection probability *σ* = 0.25, interaction strengths drawn from [−log 3, log 3], and basic reproduction numbers randomly drawn from [1.5, 2] (A). Estimated networks were obtained using the regularized Ising model from a dataset with 10,000 individuals in a stratified analysis among low-risk individuals only (B), high-risk individuals only (C), among a representative sample of the total population without correction (D), or with correction by augmenting the network with an extra node R, indicating membership to either sub-population (E). The filtered network (F) is obtained from (E) by omitting node R and the corresponding edges. Strength and type of interactions (mutualistic versus competitive) are indicated by the thickness and colour (green versus red) of edges, respectively.

Network inference regained good performance if host heterogeneity was corrected for by performing stratified analyses based on the risk variable indicating membership to either sub-population (compare purple and pink dots to black lines in Fig 4; compare B and C to A in Fig 5). At moderate sample size (1.000 and 10.000), performance was somewhat better in subgroup analyses on high-risk individuals, likely due to increased statistical power at higher prevalence of infection. The alternative correction approach by adding the risk variable as an extra node to the network, representing elevated infection risk, also performed well (compare red to turquoise in Fig 4). Graphically, the risk variable can be represented as a central node equally connected to all other nodes representing carriage of strains (Fig 5E). This dependency illustrates that elevated infection risk is associated with positivity for each strain. After filtering out the risk variable, the strain-specific interaction network is retained (Fig 5F)

## Discussion

In this paper, we evaluated whether heterogeneous pathogen strain interactions can be recovered from cross-sectional surveys tracking co-occurrence of multi-strain pathogens via statistical network inference. Using simulated data, we demonstrated unbiased estimation of pairwise interactions between multiple infectious strains, appropriately correcting for indirect interactions through other strains. Performance of network inference was shown to be strongly influenced by host heterogeneity, but we also demonstrated how this can be overcome by correcting for individual risk factors common to all pathogen strains.

Conceivably, the nature and strength of multi-strain interactions may indeed be heterogeneous, as these are modulated by within-host niche differentiation. Such differentiation may arise from slight differences in host-cell tropism, as e.g. described for oncogenic HPV infection, where genotypes of alpha-7 and alpha-9 species have distinct preference for glandular and squamous epithelium, respectively [4, 27]. Moreover, if multi-strain interactions are mediated by within-host immune responses, as hypothesized for different pneumococcal serotypes, strength of interaction may depend on antigenic overlap between strains [3]. To uncover heterogeneous interactions, it is essential to analyse datasets where the exact combination of strains in each host is recorded. Previous endeavours to infer competitive interactions from such genotype combination data found that they only performed satisfactorily in datasets where competitive interactions are particularly strong, and that accounting for host behavioural heterogeneity is more important than adding additional information via the genotype combinations [20, 21]. The importance of accounting for host heterogeneity is reiterated in our analysis, but we also demonstrate that network inference is well suited to detect small interactions at large enough sample size, provided that the mechanisms of interaction can be inferred from conditional independence structures.

Our results were obtained under a particular epidemiological model and under a steady state assumption, with between-strain interactions modelled in acquisition and clearance. We have previously shown that the odds ratio of co-occurrence in a two-strain version of this model is an exact estimator of the composite of interaction parameters defined accordingly [28]. The results of the present study should be viewed as an extension of that result to systems with an arbitrary number of strains. Network models are well suited to capture conditional independence between multiple strains, providing a clear advantage over marginal estimation of pairwise interactions as these may be confounded through indirect interactions with other strains. While we showed excellent performance of the network inference methods in this respect, we cannot ensure good performance under epidemiological models other than the one we considered. For instance, if strains interact through natural cross-immunity that is long-lasting, infections with different strains may be positively associated (i.e. as given by odds ratios greater than one) and co-occurrence patterns of current infection are no longer informative of their interaction structure [28]. Likewise, strains may interact through modification of viral or bacterial load during co-colonization, rather than through modification of acquisition or clearance. Although such interactions are better captured by records of viral or bacterial load, they are likely correlated with interactions in acquisition and clearance. Hence, network inference based on co-occurrence may retain good performance even in such situations. More extensive simulations are needed to demonstrate the robustness of the network inference methods to variations in model specification. In general, more complex mechanisms of interactions may be better identifiable from individual- or population-level longitudinal studies than cross-sectional surveys [29, 30].

Likewise, performance of network inference may depend on the way that interactions between multiple strains combine during co-infection. Previously, we demonstrated that predictors of strain-replacement in the wake of vaccination against a subset of pathogen strains, perform best when interactions in acquisition and clearance combine multiplicatively [8]. We envisage that reconstruction of heterogeneous strain interaction networks from co-occurrence data becomes more complicated if interactions combine otherwise than multiplicatively, e.g. if strength of interaction is not affected by multiplicity of infection. It should be noted that estimates obtained under GEE or Ising models are also marginal in this sense, as they only consider pairwise interactions and leave possible higher-order interactions unspecified. Conversely, graphical log-linear models implicitly incorporate a three-way interaction whenever three strains are graphically connected, allowing pairwise interactions to be modulated by a third strain [25]. Suitability of either approach is determined by the likelihood of higher-order interactions being present in a particular system, and by the quality and resolution of the available data.

While we did not directly compare the methods applied here to previous approaches, we envisage that the comparatively good performance of statistical network inference derives from their rigour and computational efficiency. These methods bypass specification of explicit models for interacting strain dynamics, and instead capitalize on the implicit relation between the interaction parameters and the expected proportions of the host population that carry different combinations of infectious strains.

Consistency of methods depends on whether the underlying interaction mechanisms are appropriately captured, but the same applies to explicit epidemiological modelling of co-infections. Both approaches may suffer from model misspecification, but if approximations hold, statistical network inference provides a particularly powerful tool, as shown here.

All network modelling approaches considered in this paper show unbiased estimation of interaction parameters, evidenced by diminishing RMSD for increasing sample size. However, we found that different methods excel in different aspects of estimation, with a general trade-off between non-discovery and false discovery. Network structure was only captured appropriately when applying some form of regularization (as in the Ising model) or penalizing for sample size (as in BIC-based log-linear model selection). In comparison, GEE or AIC-based model selection suffered from poor specificity, yielding comparatively low PPV at large sample size. Even so, these methods exhibited negligible RMSD, demonstrating that false discovery at large sample size predominantly pertains to supposedly small interaction strengths. Penalization ensures that very weak connections are set to zero; hence, the appropriateness of using AIC- or BIC-based model selection depends on the sparsity of the true interaction network being analysed. In our simulations, sparsity of the networks used to generate co-occurrence data was controlled by fixing the rate of true zeroes in the network. The extent to which true independence holds in reality should determine which class of network inference methods is to be preferred [31]. From a practical point of view, however, the important interactions were almost always correctly identified, also at limited sample size.

Finally, we show that a regression framework familiar to most epidemiologists, i.e. GEE, performs satisfactorily in estimating pairwise interactions. The GEE regression framework has the additional benefit that it offers a flexible way of separating individual risk factors common to all strains, from interactions between any two strains [23]. This framework has been employed before to study clustering patterns of HPV genotypes across risk populations, with correction for known risk factors [18]. While correction for individual-level characteristics is also possible in other methods, as demonstrated in the present study, most other graphical modelling approaches rely on the inclusion of risk factors as binary nodes to the network. This clearly limits the practical ability to correct for various sources of host heterogeneity, that are more naturally accommodated in a regression framework [26, 32]. Compared to other graphical modelling approaches, the GEE framework is not scalable to situations where the number of strains exceeds the number of observations. However, for most epidemiological studies this does not pose a real problem.

To summarize, we have demonstrated how interactions between multiple pathogen strains might be estimated from cross-sectional surveys, detecting the presence of multiple infectious strains at once. We illustrated this by applying statistical network models, that properly account for conditional independence between strains and are able to efficiently and consistently estimate heterogeneous interactions. Our work demonstrates how widely available multivariate methods may be used to identify between-strain interactions from readily available data.

## Materials and methods

### Multi-strain epidemiological model

The epidemiological model we used for simulation was previously introduced to assess the outcome of vaccination against a subset of pathogen strains present in a host population [8]. In short, this model allows for an arbitrary number of infectious strains to circulate, assuming Susceptible-Infected-Susceptible (SIS) dynamics with regard to each of the individual strains. It applies to pathogens for which naturally acquired immunity is limited so that reinfection is possible [33], as for instance could be the case for infection with human papillomavirus or with *Streptococcus pneumoniae*. For a pathogen with *n* strains, the model state space 𝒮 consists of 2*n* infection states, each denoted by the set of strains the host is infected with. See S3 Fig for a schematic representation of the possible infection states with *n* = 3. A system of ordinary differential equations describes how the proportions of individuals in each of the model states evolve over time. The equation for the change in prevalence *N*_*X*_ of any state *X* ∈ 𝒮 is generally given by

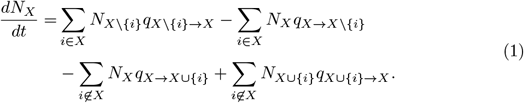

The four groups of terms on the right-hand side, moving from left to right, correspond to flows of individuals into or out of state *X* due to 1) acquisition of a strain *i* ∈ *X*, 2) clearance of a strain *i* ∈ *X*, 3) acquisition of a strain *i* ∉ *X*, and 4) clearance of a strain *i* ∉ *X*, respectively. Each term is a product of the proportion of the population in a particular state and the corresponding acquisition or clearance hazards (per capita rates) denoted by *q*. In an example with three strains and *X* = {1}, the corresponding terms are 1) *N*_ø_*q*_ø→{1}_, 2) *N*_{1}_*q*_{1}→ø_, 3) *N*_{1}_*q*_{1}→{1,2}_ + *N*_{1}_*q*_{1}→{1,3}_, and 4) *N*_{1,2}_*q*_{1,2}→{1}_ + *N*_{1,3}_*q*_{1,3}→{1}_.

The baseline hazard of clearance for an individual only infected with strain *i* is denoted by *q*_{*i*}→ø_ and the baseline hazard for a completely susceptible individual to acquire strain *i* is given by

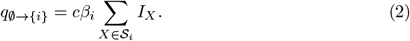

Here, *c* is the per capita rate at which hosts make contacts that are relevant for pathogen spread, *β*_*i*_ is the probability of successfully acquiring strain *i* given contact with an individual infected with strain *i*, and 𝒮_*i*_ ⊂ 𝒮 is the subset of states containing strain *i*.

Potential interactions between strains are modelled through modification of the baseline acquisition or clearance hazards. The model allows for different structures of interactions, but we only consider the pairwise-symmetric multiplicative structure (see [8] for the alternative structures). To be exact, we assume each strain that is carried to contribute multiplicatively to the hazard of acquiring (or clearing) the incoming (or outgoing) strain. Hence, the hazards of acquisition and clearance in the presence of other strains are

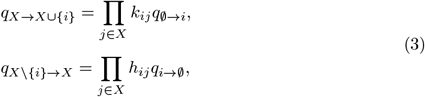

where the involved interaction parameters are pairwise-symmetric, i.e. *k*_*ji*_ = *k*_*ij*_ and *h*_*ji*_ = *h*_*ij*_ for all *i* and *j*. Defined as such, *k*_*ij*_ and *h*_*ij*_ are essentially hazard ratios that act on the baseline hazards. In particular, values equal to one imply no interaction and the deviation from one indicates the strength of interaction. For the mode of acquisition, *k*_*ij*_ *<* 1 and *k*_*ij*_ *>* 1 indicate competitive and mutualistic interactions, respectively. Conversely, for the mode of clearance, *h*_*ij*_ *<* 1 and *h*_*ij*_ *>* 1 indicate mutualistic and competitive interactions, respectively. Previously, we have shown that when interaction is present in both modes, the overall interaction could be summarized by the ratio of the two interaction parameters *k*_*ij*_*/h*_*ij*_, with *k*_*ij*_*/h*_*ij*_ *<* 1 and *k*_*ij*_*/h*_*ij*_ *>* 1 indicating competitive and mutualistic interactions, respectively.

Finally, the basic reproduction number of each strain *i* is defined by *R*_0,*i*_ ≡ *cβ*_*i*_*/q*_{*i*}→ø_. In the absence of interactions, only strains having *R*_0,*i*_ *>* 1 would be expected to survive, with marginal prevalence given by 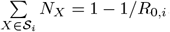 Note, however, that *R*_0,*i*_ *>* 1 is neither sufficient nor required for survival in the presence of strain interactions, and that rank marginal prevalence need not reflect rank reproduction number. For simplicity, we fixed *c* = 3, set *q*_{*i*}→ø_ = 1 for all strains, and drew *β*_*i*_ randomly to obtain random *R*_0,*i*_.

### Network construction

In the interaction networks we consider in this paper, nodes represent pathogen strains and edges the presence of a pairwise interaction between two strains. By connecting each pair of strains with fixed connection probability *σ*, independent from other edges, the strain interaction networks resemble Erdös–Rènyi random graphs that become more saturated with higher connection probability [34].

Binary networks only indicating presence or absence of interactions were converted to weighted networks by attributing strengths of interaction *x*_*ij*_, drawn uniformly between [−*θ, θ*], with *θ >* 0 denoting the maximum strength of interaction. In settings where we considered interaction in acquisition only, exp(*x*_*ij*_) was used as *k*_*ij*_ in the epidemiological model. When we also considered interaction in clearance, aside from *x*_*ij*_, which we drew uniformly from [−*θ, θ*], we also drew an additional random number 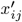 uniformly from [−*θ/*2, *θ/*2]. 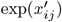 was used as *k*_*ij*_ and 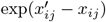 as *h*_*ij*_.

### Sampling cross-sectional datasets

We sampled random cross-sectional datasets [*Y*_1_, *Y*_2_, …, *Y*_*m*_]^*T*^ from a multivariate binomial distribution that coincides with the steady state of the described epidemiological model, where *m* is the number of i.i.d. observations, i.e. individuals. For each individual *l, Y*_*l*_ = [*Y*_1,*l*_, *Y*_2,*l*_, …, *Y*_*n,l*_] denotes the sequence of Bernoulli random variables [35], with *Y*_*i,l*_ denoting the presence of strain *i* in individual *l*. It follows that *P* (*Y*_*i,l*_ = 1 ∀*i* ∈ *X, Y*_*j,l*_ = 0 ∀*j* ∉ *X*) = *N*_*X*_, where *N*_*X*_ is the steady state solution of model (1).

### Statistical network inference

In previous work, we and others have shown that in a two-strain version of model (1), the prevalence of being simultaneously infected with two strains that do not interact, corresponds to the product of marginal prevalence of both strains, provided that host death in age groups susceptible to infection is negligible [21], and there are no unobserved common risk factors [28]. Moreover, the odds ratio of co-occurrence in the two-strain model corresponds to the ratio of interaction parameters in acquisition and clearance *k*_12_*/h*_12_ [28]. When there are more than two strains, however, the correspondence between pairwise association measures and corresponding pairwise composite interaction parameters *k*_*ij*_*/h*_*ij*_ might no longer hold, and deviation from independence between two non-interacting strains could be induced by a third strain that interacts with both. The objective here is to capture these (possibly heterogeneous) pairwise interaction parameters from cross-sectional prevalence surveys for more than two strains. For this purpose, we make use of various network inference methods (S1 Appendix).

The first statistical network inference method we applied are graphical modelling approaches based on log-linear analysis [36]. This technique is generally used to examine the relationship between more than two categorical variables, here denoting presence-absence of each pathogen strain in a host population. To reconstruct networks of strain-specific interactions from co-occurrence data with up to 10 circulating strains, we searched through the subset of decomposable log-linear models, i.e. models whose dependence graph is triangulated [25]. Selection was applied both in a forward (focussed on adding edges) and backward (focussed on removing edges) fashion, using AIC or BIC as selection criterion. Strengths of interaction were quantified a posteriori by calculating odds ratios using conditional maximum likelihood estimation from the contingency tables implied by the selected model.

Alternatively, we used the Ising model to reconstruct the simulated networks [24, 37]. Here, the probability of a certain pathogen strain being present is modelled as a function of other strains being present at the same time. The Ising model can be shown to be equivalent to certain kinds of log-linear models, but interactions are at most pairwise and the dependence graph does not need to be triangulated. To estimate presence and strength of interactions, we made use of regularized logistic regressions: iteratively, one variable is regressed onto all others, with a penalty imposed on the regression coefficients to obtain a sparse network representation [38], and with model selection based on the extended BIC [24].

Lastly, we used generalized estimating equations (GEE) to estimate pairwise odds ratios between all strains under consideration. In modelling the associations between multiple strains concomitantly, we used the alternating logistic regression algorithm of the GENMOD procedure in SAS statistical software [26, 32]. The regression framework facilitates assessment of strain-specific interactions on the basis of Wald tests. To correct for multiple hypothesis testing, we made use of the Benjamini–Hochberg procedure [39].

### Host heterogeneity

In the sensitivity analysis with host heterogeneity we considered a version of the described multi-strain epidemiological model with two sub-populations that differ in their contact rates relevant for pathogen spread. The proportions of hosts in the high- and low-contact sub-populations are denoted by *p*_1_ = 20% and *p*_2_ = 80%, and the corresponding contact rates by *c*_1_ and *c*_2_, respectively. The values of *c*_1_ and *c*_2_ were chosen to obtain a coefficient of variation of 80%. Contacts between the two sub-populations follow a classical pattern based on assortativity fraction *ϕ* within host types, that takes the value 0 when mixing between hosts is random and 1 when fully assortative [33]. Here, we fixed *ϕ* = 50%. The contact rate *c*_*sr*_ between a ‘transmitting’ individual from sub-population *s* and a ‘receiving’ individual from sub-population *r* is given by (see S2 Appendix for derivation)

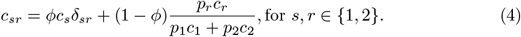

Here, *δ*_*sr*_ is the Kronecker delta (with *δ*_*sr*_ = 1 if *s* = *r* and zero otherwise). Defined accordingly, the baseline hazard for a susceptible individual in sub-population *r* to acquire strain *i* is given by

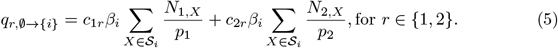

Here, *N*_*r,X*_ is the proportion of individuals in sub-population *r* ∈ 1, 2 and infection state *X*, with 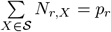 Differential equations for *N*_*r,X*_ were defined analogously as in model (1) and the hazard for acquisition in presence of infection with other strains as in equation (3), e.g.

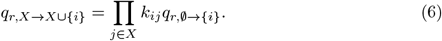

For network reconstruction in the stratified analysis, we performed analyses separately within either sub-population. For the alternative analysis with correction, we included an additional variable R indicating to which sub-population each individual belongs. In the GEE regression framework, we included R as an explanatory variable (irrespective strain). In the other network approaches, we added R to the sequence of variables denoting strain-specific presence, i.e. 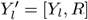. In effect, the conditional independence structure inferred from 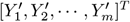 is augmented by one node representing elevated infection risk, next to nodes representing strains.

## Supporting information

**S1 Fig. Schematic representation of the data generation process**. For simplicity, the workflow is illustrated for a system with *n* = 3 strains but the same principle was applied in base-case and sensitivity analyses for systems of *n* = 5 or *n* = 10 strains.

**S2 Fig. Scatter plots of the estimated against the true interaction parameters**. Four scatter plots of the estimated against the true interaction parameters under GEE or Ising models, with or without correction for contact rate.

**S3 Fig. Schematic representation of the epidemiological model with** *n* = 3 **strains**. The eight (2^*n*^) infection states and transitions pertaining to acquisitions. For convenience, the reverse transitions (i.e. clearances) are not shown, and infectious states are not shown in set notation with brackets, unlike the formulas in the main text.

**S1 Table. Performance measures of statistical network inference in the sensitivity analysis with host heterogeneity**. In the base-case analysis, interactions between strains were generated with connection probability *σ* = 0.25, interaction strengths indicating interaction in acquisition drawn from [− log 3, log 3], and basic reproduction numbers randomly drawn from [1.5, 2]. In this sensitivity analysis, the average contact rate relevant for pathogen spread is the same as in the base-case simulations, but 80% of hosts is assumed to have below-average contacts and 20% above-average contact. Mixing between sub-populations occurred pseudo-assortatively with an assortment fraction of 50%. Abbreviations: GEE, generalized estimating equations; PPV, positive predictive value; RMSE, root-mean-square error.

**S1 Appendix. Statistical network inference**.

**S2 Appendix. Host heterogeneity**.

